# A novel Energy Landscape method incorporating the Latent Dirichlet Allocation topic model and the pairwise Maximum Entropy model, revealing the significant contribution of Bactericides to the development of Inflammatory Bowel Disease

**DOI:** 10.1101/2023.04.19.537426

**Authors:** Kaiyang Zhang, Shinji Nakaoka

**Affiliations:** Graduate school of Life Science, Hokkaido University; Faculty of Advanced Life Science, Hokkaido University

## Abstract

The dysbiosis of microbiota has been reported to be associated with numerous human pathophysiological processes, including Inflammatory Bowel Disease (IBD). With advancements in highthroughput sequencing, various methods have been developed to study the alteration of microbiota in the development and progression of diseases. However, a suitable approach to assess the global stability of the microbiota in disease states through time-series microbiome data is yet to be established. In this study, we introduce a novel Energy Landscape construction method, which incorporates the Latent Dirichlet Allocation (LDA) model and the pairwise Maximum Entropy (MaxEnt) model, and demonstrate its utility by applying it to an IBD time-series dataset. Through this method, we obtained the “energy” profile for the potential patterns of microbiota to occur under disease, indicating their stability and prevalence. The results suggest the potential contribution of several microbial genera, including *Bacteroides, Alistipes*, and *Faecalibacterium*, as well as their interactions, to the development of IBD. Our proposed method provides a novel and insightful tool for understanding the alteration and stability of the microbiota under disease states and offers a more holistic view of its complex dynamics at play in microbiota-mediated disease.

## 2 Introduction

The microbiota residing in the human body plays a crucial role in maintaining health and wellbeing, with varying composition in different body sites, including the mouth, vagina, skin, and particularly, the intestinal tract [1]. It has also been dubbed as “forgotten organ” due to its collective and complex metabolic activity [2]. Bowel dysbiosis, an imbalance in the composition of the microbiota, has been linked to numerous diseases including gastrointestinal disorders such as inflammatory bowel disease (IBD), irritable bowel syndrome (IBS), type 2 diabetes (T2D), and colorectal cancer (CRC); systemic diseases such as metabolic disorders in obesity; and even central nervous system (CNS) disorders like autism spectrum disorder (ASD) [3]. A comprehensive understanding of the impact and mechanisms of microorganism-host interactions is essential for the diagnosis and treatment of associated diseases.

Recent advancements in high-throughput Next Generation Sequencing (NGS) techniques, such as 16S rRNA sequencing and metagenomic sequencing, have greatly improved our ability to profile the microbiota. After that, analysis of those obtained microbial NGS data is becoming a new challenge in investigating of microorganism–host interactions and several methods have been developed to analyze microbial NGS data. However, previous studies tend to consider the relationship between microbiota and disease from the perspective of “one microbe-one disease”, which is an oversimplification and may fail to explain the complex mechanism [1]. Instead, the focus should be on the comprehensive dysbiosis of the microbiota, including qualitative and quantitative changes, metabolic activities, and local distribution [4]. Longitudinal studies that collect time-series microbiome data can mitigate sampling biases and cover sufficient periods of disease development in participants, providing deeper insights into the dynamic alterations and interactions of the microbiota components in dysbiosis and disease pathogenesis. To this end, we present an Energy Landscape construction approach that combines two unsupervised machine learning models: Latent Dirichlet Allocation (LDA) and pairwise Maximum Entropy (MaxEnt), to obtain insight into global stability of the microbial community from longitudinal time-series data.

The Latent Dirichlet Allocation (LDA) model is a widely applied unsupervised machine learning method in natural language processing (NLP). It models text through a three-level hierarchical Bayesian model, with ‘topic-word’ and ‘document-topic’ multinomial distributions and a Dirichlet prior.[5]. In the context of microbial abundance profiles, the LDA model can identify ‘microbial assemblages’ by grouping taxa according to their co-occurrence features [6, 7], similar to the ‘topics’ in NLP studies. The pairwise MaxEnt model, for another thing, provides a second-order maximum entropy model that captures a single node’s firing rates and the pairwise interactions in biological system, assuming no higher-order interactions[8, 9]. This model has been demonstrated to accurately describe neural systems using time-series MRI data[10, 8]. In our Energy Landscape construction approach, the microbial assemblages are modeled as having pairwise interactions, akin to neuronal activity, and can be analyzed using the pairwise MaxEnt model. The LDA model will cluster the microbial abundance profile into microbial assemblages, while the pairwise MaxEnt model will construct an ‘energy’ profile for all potential activity patterns of microbial assemblages, from which we can determine the occurrence patterns of assemblages in specific health conditions. In this study, we applied the Energy Landscape with these two models to 16S rDNA time-series data from the HMP2 IBD database. Our evaluation of the obtained energy profile showed that some microbial assemblage patterns and related “energy basins” might be recognized as globally stable under health conditions including non-IBD, Ulcerative Colitis (UC), and Crohn’s Disease (CD). Interestingly, our results indicate that the patterns, characterized by the sole activation of various levels of the genus *Bacteroides*, are present in the stable patterns of CD and imply the contribution of *Bacteroides* to the deviation of microbiota in the development of IBD. Moreover, the presence of other members in these activated assemblages in stable patterns highlights the potential interaction and joint effect of multiple genera in the dysbiosis of microbiota in IBD development.

This novel method offers a fresh perspective into the microbiota by providing a quantitative description of a dynamic biological system and depicting its global stability. It may help researchers better understand the microorganism-host interaction and its significance for human health.

## 3 Materials and Methods

### 3.1 Metagenomic time-series dataset

We use the dataset from the Onset of Inflammatory Bowel Disease (IBD) of The Integrative Human Microbiome Project (iHMP)(NIDDK U54DE023798)[11]. The dataset contains taxonomic profiles of fecal samples’ 16S rDNA sequencing from the project’s participants. Those taxonomic profiles of each participant were collected repeatedly over a study period. Here, we choose each participant’s first successive ten time-series samples to process, and filtered the participants with less than ten samples. Finally, the sample size comprises 1300 samples collected from 130 participants with each participant contributing 10 samples. Several participants have been diagnosed with two major types of IBD: Crohn’s Disease (CD), Ulcerative Colitis (UC), while the remaning participants without IBD (non-IBD) serve as control.

### 3.2 The sample size

The table 1 presents the classes in this study. In the LDA modeling step, only the first sample from each participant is used as the input (*N* = 130) to avoid the bias resulting from the homogeneity of composition from the same participant. After we have acquired the LDA model with the parameter ***φ***_*i*_, the model is applied to the 780 samples (as detailed below) as the next phase’s input. During in the pairwise MaxEnt modeling phase, the modeling is conducted separately for three disease types. In order to facilitate comparison, balanced input classes (*N* = 26 × 10 = 260), consisting 780 samples in total, are chosen for for each modeling execution (CD,UC,non-IBD).

**Table 1:** The classes in the study. The numbers without brackets represent the number of participants, and in bracket represent the number of samples.

|  | Disease types |  |  |
| --- | --- | --- | --- |
| Modeling steps | CD | UC | non-IBD |
| LDA modeling | 65(65) | 38(38) | 27(27) |
| pairwise MaxEnt modeling | 26(260) | 26(260) | 26(260) |

### 3.3 Latent Dirichlet Allocation(LDA) modeling

In accordance with the principle of the LDA model, the microbiota, or microbial community, is comprised of a series of single “occurrence-event”(hereinafter referred to as occurrence). An occurrence is defined as the solitary presence of a taxonomic unit. Each occurrence belongs to a latent attribute: microbial assemblage. The probability profile of the taxonomic unit characterizes the likelihood of the microbial assemblage’s presence, and the candidate taxa can be considered as the constituents of the assemblage, each with varying weights. Hence the imaginary generation process of a microbial community can be assumed as follows: Initially, under specific distribution, an assemblage of one occurrence is assigned; Subsequently, under another specific distribution, determined by the assigned assemblage, the taxon of the occurrence is assigned. This process repeats, and ultimately, the occurrences combine to form the microbial community.

If *I* microbial assemblages and taxon level of the genus in *N* microbial samples are decided, the assemblages in *N* samples follow the multinomial distributions with parameters ***θ*** _*n,n*∈ (1,…,*N*)_ ; while the genera in *I* assemblages follow multinomial distributions with parameters ***φ***_*i,i* ∈ (1,…,*I*)_. The vector 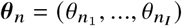 represents the probability of the occurrences of *I* microbial assemblages in the *n*th sample, and the vector 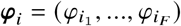 represents the probability of the occurrences of *F* genera in the *i*-th assemblage. Thus we can regard ***θ*** and ***φ*** as the “abundance” of assemblages in a specific sample and the weight of genera in one specific assemblage, respectively.

After parameter estimation of ***θ*** _*n*_ and ***φ***_*i*_ are completed, the *F*-dimension original genera abundance profile is reduced to *I*-dimension assemblages abundance profile, which is feasible for the following Maximum Entropy Modeling.

Here we choose *I* = 9 to get the number of assemblages that reduce the computational cost of pairwise Maximum Entropy modeling and keep the interpretability (see also discussion). The LDA modeling is conducted by sklearn.decomposition.LatentDirichletAllocation package[12, 13, 14].

### 3.4 pairwise Maximum Entropy(MaxEnt) Modeling

We fit the pairwise MaxEnt model according to the manners in its previous applications for neuroscience [9, 10, 15]. In the pairwise MaxEnt model, the goal is to maximize the information entropy of probability distribution under the Maximum Entropy Principle, and adjust the model to match correlations of strength of individual assemblage and all assemblage pair interactions from observation data, which represented by the constraints of ⟨***σ***_*i*_⟩ and ⟨***σ***_*i*_***σ*** _*j*_ ⟩. ⟨***σ***_*i*_⟩ and ⟨***σ***_*i*_***σ*** _*j*_ ⟩ are defined as following:

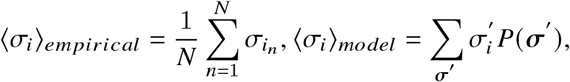

where 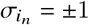 is the occurrence state of the *i*-th assemblage on the *n*-th sample;

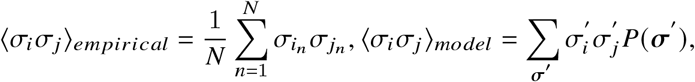

where *i* and *j* represent two different assemblages.

Then the model can be given by:

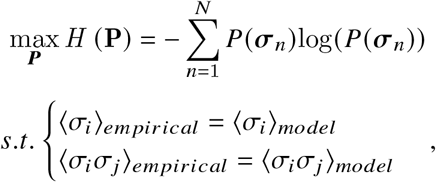

where *empirical* represents the empirical results and *model* represents the expected value given by the model, respectively.

The pairwise MaxEnt model gives the probability of assemblage patterns to occur ***σ*** in the following distribution:

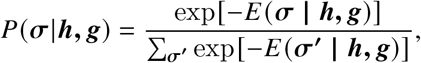

where

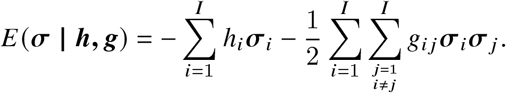

The ***h*** and ***g*** are the parameters that need to be estimated from the data, representing the tendency to the occurrence of one assemblage and the interaction between two assemblages, respectively. Positive and negative values of ***g*** are interpreted as promotional and inhibitory interactions, respectively. We estimate the parameters through the maximum-likelihood method[15]. Here we solve

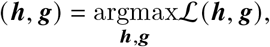

where ℒ (***h, g***) is the likelihood function given by

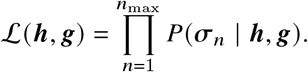

The likelihood is maximized by updating ***h*** and ***g*** in the gradient ascent scheme till convergence:

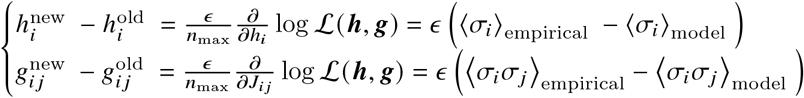

where *new* and *old* represent the values after and before a single updating step, and *ε* > 0 is a constant controlling the step size.

### 3.5 Definition of the occurrence state in assemblage pattern

In the pairwise MaxEnt model, the input assemblage pattern with binary elements is necessary. Here the assemblage pattern is defined as ***σ***, where the value of each microbial assemblage ***σ***_*i,i*∈ (1,…,*I*)_ is assigned either 1 or -1 according to the “occurrence state”. This state represents whether the specific microbial assemblage has a relatively high abundance on a sample. Recall that the assemblage’s abundance in each sample is assigned by the parameter ***θ*** _*n*_ given by the LDA model. According to the property of multinomial distribution in the LDA model, its parameter of assemblage’s probability can measure the occurrence of the assemblage and be regarded as comparable abundance in a microbial community. For example, if *i*-th assemblage of *n*-th sample has a higher probability parameter than that of *m*-th sample 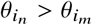, given by the LDA modeling, we consider the *i*th assemblage shows higher abundance on the microbial community of *n*-th sample than *m*-th sample.

Then, a threshold to define the “relatively high” or “activated” to binarize the assemblage’s abundance is required. Here we assign the occurrence state ***σ***_*i*_ under the following rule:

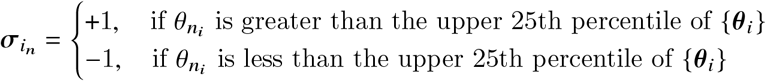

where 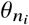 is the probability of *i*th assemblage in *n*th sample given by the LDA model,{***θ***_*i*_} the set of the probability of *i*th assemblage in all samples. The occurrence state of assemblage in a microbial community corresponds to the binary spike state of a single neuron in Schneidman’s study, which assigns the response of the neuron in a binary state of 1 (spike) and 0 (not spike)[9]. We can integrate two models through this definition by transferring the output from LDA modeling ***θ*** to the binary input for pairwise MaxEnt modeling ***σ***.

### 3.6 Energy Landscape

The distribution we obtain from pairwise MaxEnt model has the form of the Boltzmann distribution in statistical mechanics:

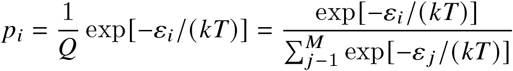

where ε_*i*_ is the energy of the system at state *i, k* the Boltzmann’s constant, *T* the temperature [9]. Recalling the distribution of assemblage pattern *P*(***σ***|***h, g***) we have obtained, we refer *E* (***σ***| ***h, g***) to the energy of the system in Boltzmann distribution.

According to the obtained parameters ***h*** and ***g*** and the function *E* (***σ***| ***h, g***), we then assign an energy value to all potential assemblage patterns. We consider that the assemblage patterns with high energy are suggested to be unstable and have a low probability of occurring and vice versa.

Once we have the energy table for all assemblage patterns, we can construct the Energy Landscape. The Energy Landscape is constructed in the manner of Ezaki’s study[15]. First, the neighbor pattern of assemblage pattern ***σ***, denoted by ***σ***′, is defined as the pattern with only a single assemblage state difference. For example, the assemblage pattern with nine assemblages ***σ*** = (1, −1, −1, −1, −1, −1, −1, −1, −1, −1) and ***σ***^′^ = (−1, −1, −1, −1, −1, −1, −1, −1, −1, −1) are neighbor patterns to each other. We assume the neighbor patterns have a close relation to the original pattern, and the pattern transition to the neighbor patterns is the first step of any further transition. Second, the energy of each pattern *E* (***σ***′) is compared to all its neighbor patterns *E* (***σ***^**′**^). If *E* (***σ***_*d*_) is the minimum in the comparison, this *d*th assemblage pattern ***σ***_*d*_ is considered as a local minimal pattern (LMP). There are several LMPs in one energy landscape, which correspond to the bottom of the basins in the energy landscape. These basins reflect the low energy locations of the system. Finally, any assemblage pattern belongs to the basin of a local minimal pattern through the path linking each pattern to its neighbor pattern with the lowest energy (see the result section). The motivation tendency of the microbial system can be interpreted as the assemblage pattern transits to its neighbor pattern with higher stability and finally towards the LMPs with locally highest stability. Overall, we can obtain the energy landscape that illustrates the energy relationship of the dynamic microbial system and especially those stable patterns which might contribute to specific health states of the host.

## 4 Results

### 4.1 LDA modeling result

Note that parameters ***θ*** and ***φ*** represented the abundance of assemblages and the composition of assemblages, respectively. The composition varied among assemblages. Some assemblages were clearly dominated by a single genus, such as assemblage #6 and #4 dominated by genus *Bacteroides* (0.89) and *Prevotella* (0.87), respectively. While two or more genera mildly dominated others: *Bacteroides* (0.52) and *Faecalibacterium* (0.22) in assemblage #1, *Akkermansia* (0.35) and *Lachnospiraceae* (0.19) in assemblage #2, *Escherichia* (0.22) and *Lachnoclostridium* (0.14) in assemblage #3, *Alistipes* (0.29) and *Faecalibacterium* (0.20) in assemblage #5, *Veillonella* (0.64) and *Fusobacterium* (0.16) in assemblage #7, *Escherichia* (0.29) and *Prevotella* (0.27) in assemblage #8, *Roseburia* (0.60) and *Haemophilus* (0.11) in assemblage #9. (Figure 2 B, Table 2) The components of one assemblage can be regarded as sharing similar characteristics and contributing to the assemblage’s effects to the host.

**Figure 1:**
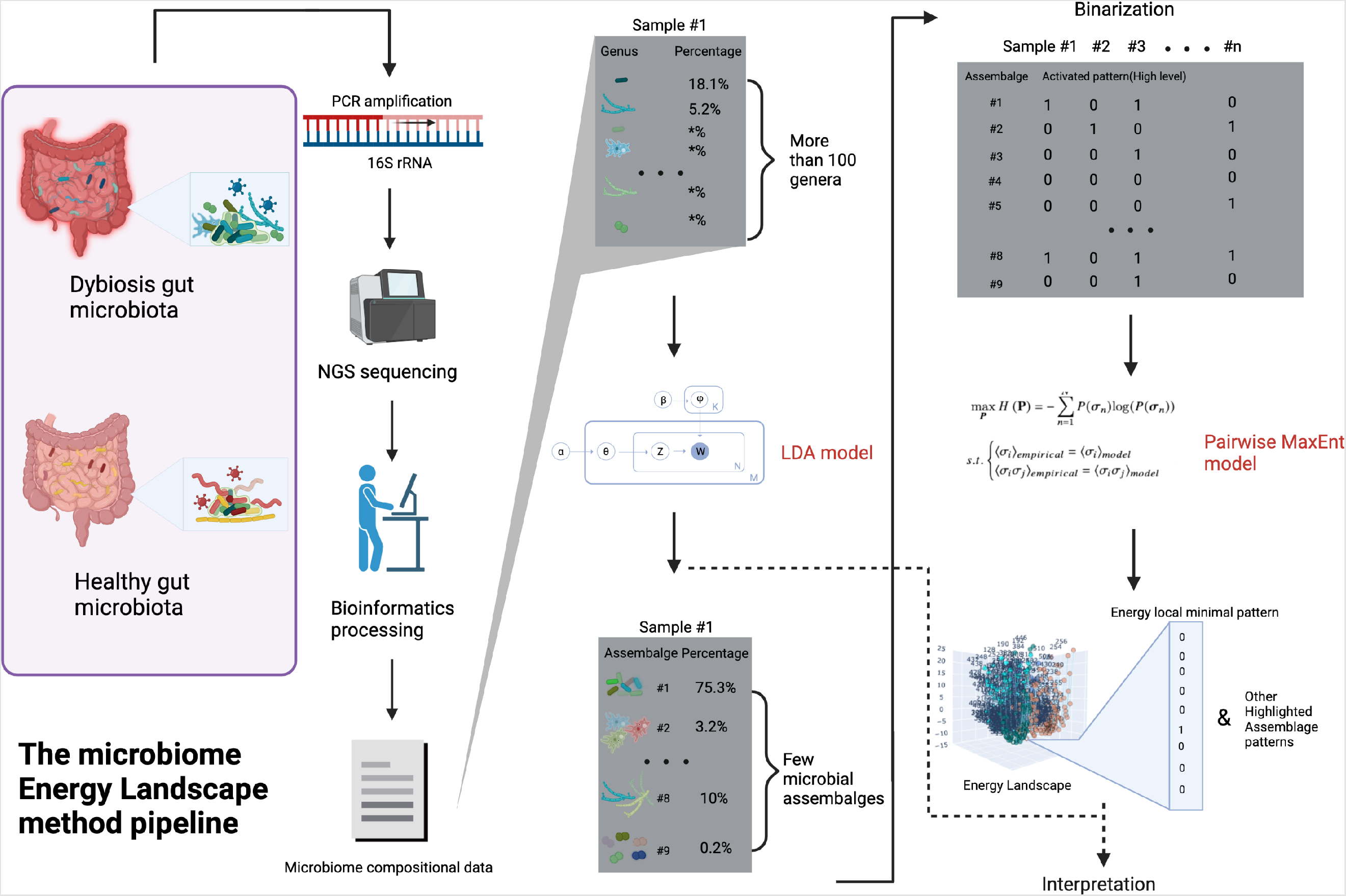
The scheme of microbiome Energy Landscape method.s

**Figure 2:**
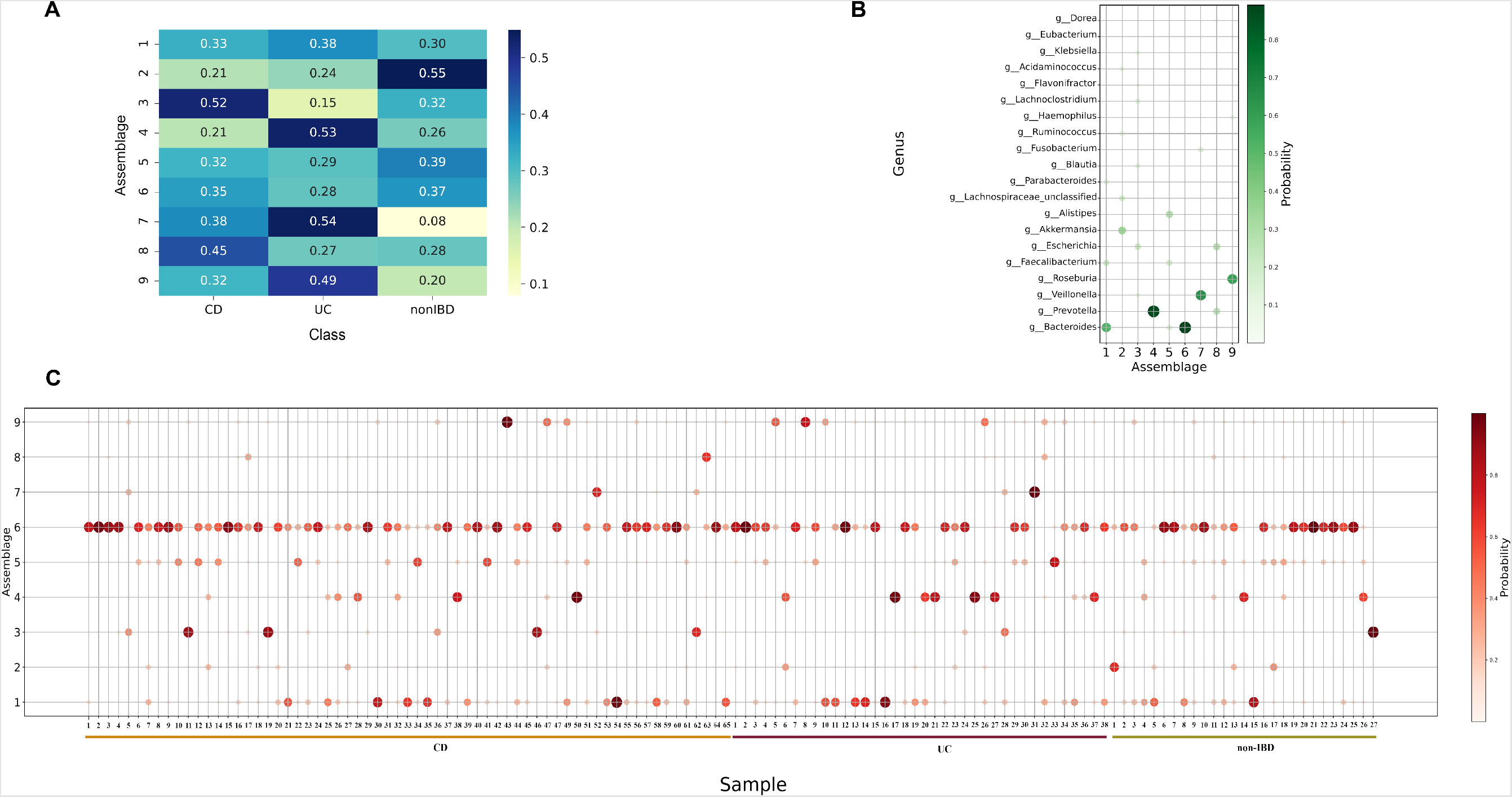
The results of LDA modeling and computation. **A**, The assemblages’ abundance among three disease classes, compared by the average probability of the assemblage in each class (***θ*** _*LDAmodeling*_)_*average*_. The values for each assemblage are scaled to one. **B**, The composition of each assembles ***φ*** given by the LDA model. Only the top ten frequent genera for each assemblage are depicted. **C**, The abundance of assembles among the 130 samples ***θ*** _*LDAmodeling*_ given by the LDA model. 9

**Table 2:** The dominant genera for assemblages. The values in brackets represent the weight of genera *φ*.

| Assemblages | Dominant genera |
| --- | --- |
| 1 | *Bacteroides* (0.52) and *Faecalibacterium* (0.22) |
| 2 | *Akkermansia* (0.35) and *Lachnospiraceae* (0.19) |
| 3 | *Escherichia* (0.22) and *Lachnoclostridium* (0.14) |
| 4 | *Prevotella* (0.87) |
| 5 | *Alistipes* (0.29) and *Faecalibacterium* (0.20) |
| 6 | *Bacteroides* (0.89) |
| 7 | *Veillonella* (0.64) and *Fusobacterium* (0.16) |
| 8 | *Escherichia* (0.29) and *Prevotella* (0.27) |
| 9 | *Roseburia* (0.60) and *Haemophilus* (0.11) |

From the abundance of assemblages in the LDA modeling samples (*N* = 130), we can see that the abundance showed a strong imbalance between assemblages. In most samples, the abundance of assemblage #6 was notably higher than that of other assemblages (Figure 2 C). Although the average abundance of assemblages showed a difference between three classes (Figure 2 A), which was calculated by 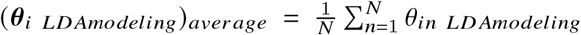 where *N* = {CD : 65, UC : 38, non-IBD : 27} only included the samples for training LDA model, there was no assemblage showing a significant difference in abundance between the classes (Data not shown). This may imply the difference was caused by extreme abundance in a few samples instead of overall different levels of the assemblage.

Among the dominant components defined by top five highest probability in ***φ***, the assemblages shared several common genera. Figure 3A shows the relation between assemblages by the common dominant genera. Note that assemblages #1, #6, and #4 share three common genera with each other and create a small relation cycle, and only assemblage #9 has no connection to other assemblages. Intuitively, the dominant genera of each assemblage shape the function of assemblages, so we might learn the functional connection from the such a relation network. Figure 3B lists all these dominant genera. The *Bacteroidess, Faecalibacterium* and *Parabacteroides* were the most frequent genera with the 6,4,3 times, respectively, and the other genera are all less than three times.

**Figure 3:**
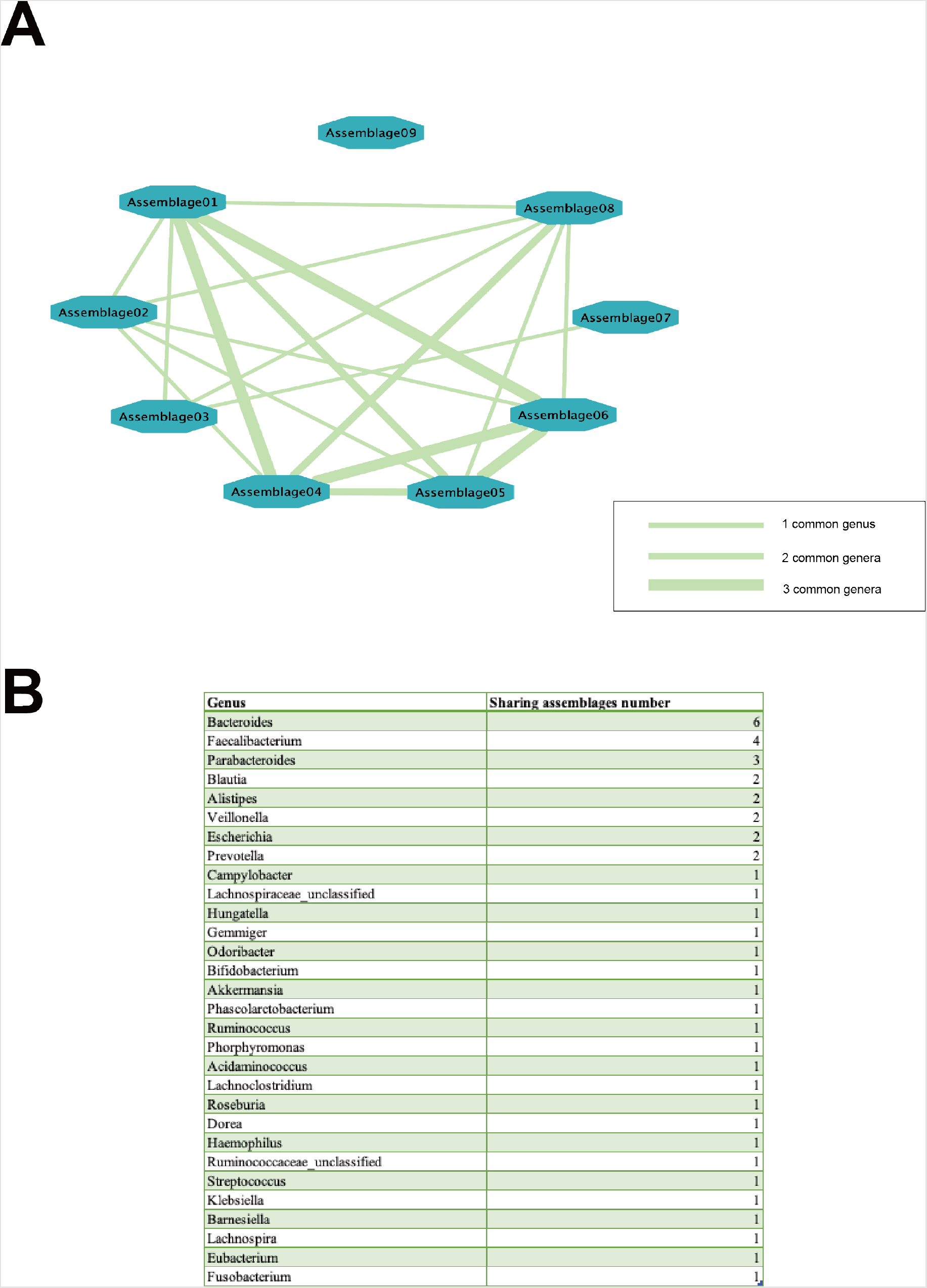
Common genera of assemblages. **A**, the network shows the relation between assemblages, where the edges are weighted by the common genera number within the top five dominant components of each assemblage. **B**, The table shows the top five dominant genera of all assemblages and the times they recur on different assemblages.

After estimating the parameters ***φ***_*i,i* ∈ (1,…,*I*)_, we applied the LDA model to the samples (*N* = 26 × 10 × 3 = 260 × 3 = 780) and obtained the weight of assemblages ***θ*** for following pairwise MaxEnt modeling.

### 4.2 Pairwise MaxEnt modeling result

The parameters ***h*** and ***g*** were obtained from the modeling of pairwise MaxEnt in three classes with the same sample size (*N* = 26 × 10 = 260). The Figure 4A shows the tendency for occurrence of single assemblages ***h***. By definition the low value of ***h*** means the low energy and high probability to occur. Notably, we can see that assemblages #6 and #1 dominated by *Bacteroides* had obviously low values than other assemblages. They had lower values in the CD class than in UC and non-IBD classes. The Figure 4B shows the pairwise interaction between assemblages. Here the high value of ***g*** in two specific assemblages means their co-occurrence contributes to low energy, and thus high probability. Several differences in interaction features among the three classes can be observed. Interestingly, the value of interaction between assemblages #1 and #6 had clearly a low value in non-IBD class compared with CD and UC classes.

**Figure 4:**
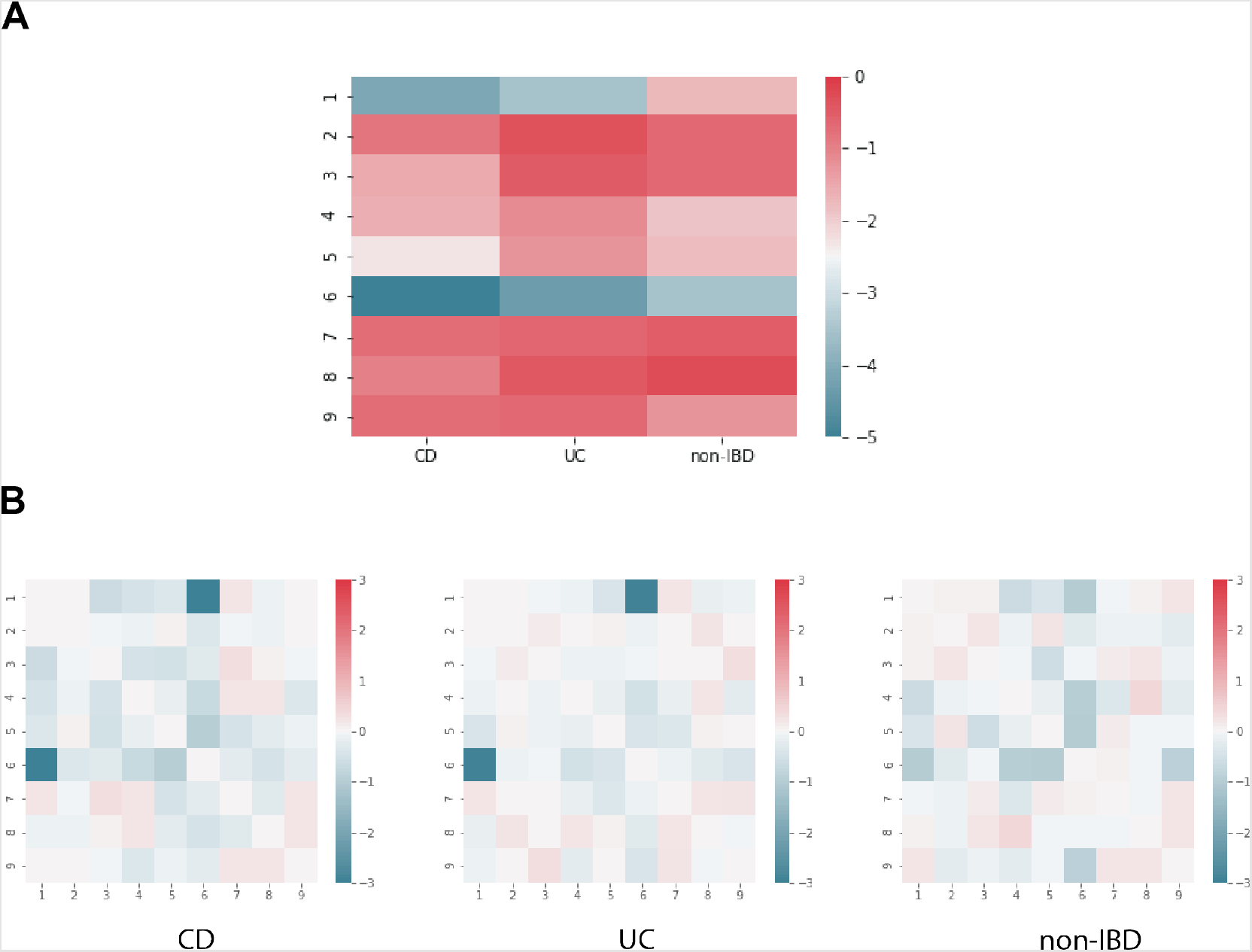
Pairwise MaxEnt results for three classes. **A**, the heatmap shows the tendency for occurrence of single assemblages ***h*** obtained from the pairwise MaxEnt model. **B**, three heatmaps describe pairwise interactions between assemblages ***g*** obtained from the pairwise MaxEnt model.

Besides, the modeling results of other CD classes with different participants showed similar features on both parameters (Additional Figure 1 A,B), which would support the reproducibility of our method.

### 4.3 Energy landscape

The energy landscape was constructed through the energy of 512 assemblage patterns given by the energy function with parameters ***h*** and ***g*** in the method section. Figures 5A, B, and C depict the energy of assemblage patterns in three classes, respectively, by drawing the line plot linking the patterns with their steepest energy descent neighbor pattern (see the method section with energy value as Z-axis). Figure 5D shows the LMPs of each class, which represent the assemblage patterns with locally low energy and high stability. Four LMPs were observed in the CD class: pattern #2, pattern #17, pattern #33, pattern #137; while two LMPs were observed in UC and non-IBD classes, respectively: pattern #33 and pattern #455 in UC, pattern #9 and pattern #33 in non-IBD. Within these LMPs, pattern #2, #17, #33 and #9 had only a single positive assemblage, while patterns #137 and #455 had multiple positive assemblages. Notably, the pattern #33 was shared in all three classes, and other patterns were unique in specific class.

**Figure 5:**
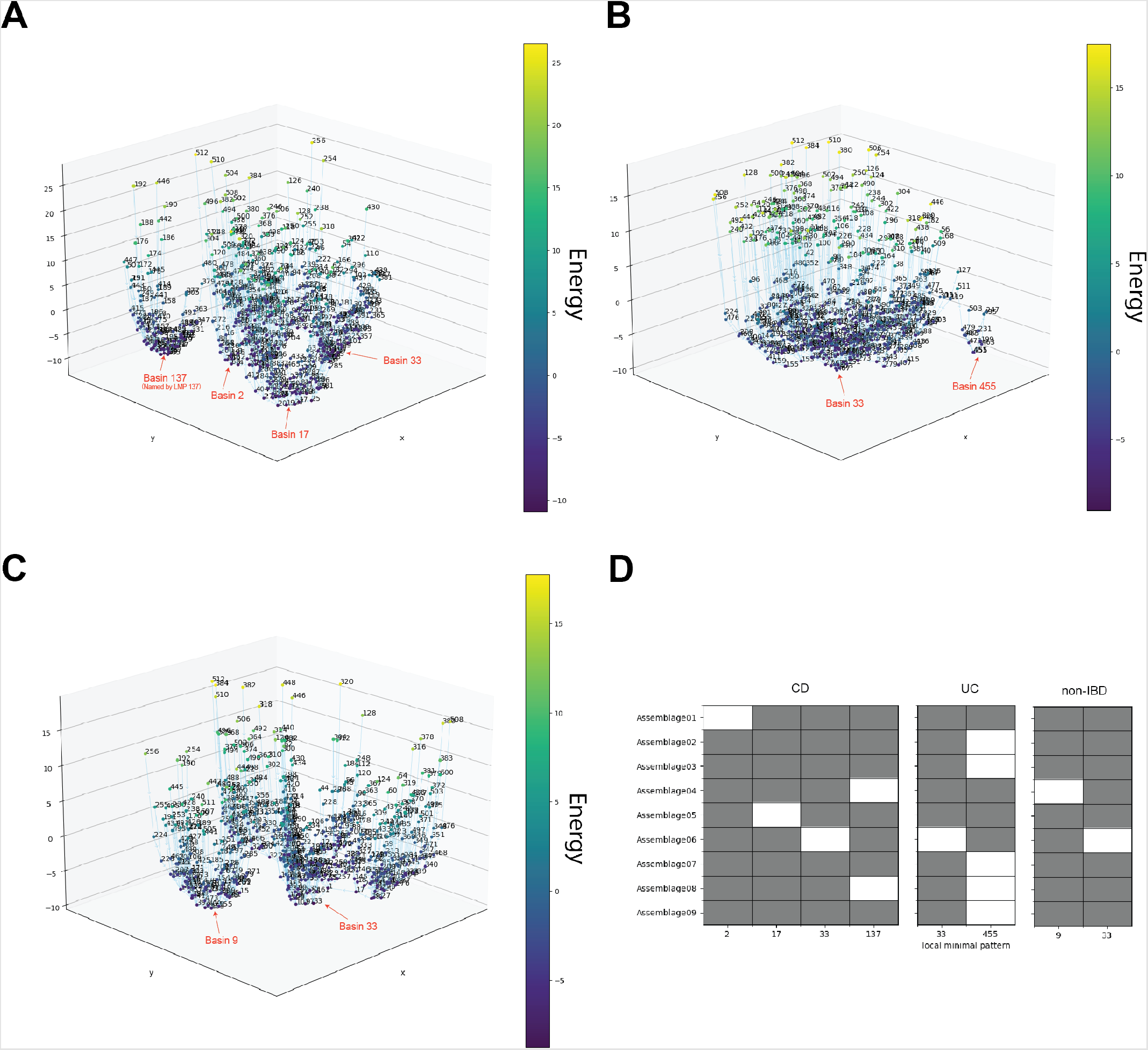
Energy landscape constructed according to the pairwise MaxEnt modeling results. **A, B, C**, the 3D line plot showing the energy of all the patterns of 9 assemblages in CD, UC, and non-IBD class, respectively. Each assemblage pattern is connected to its neighbor pattern with the steepest energy descent or to itself when it is a local minimal pattern. **D**, the composition of each LMP in three classes: CD, UC, non-IBD, from left to right, respectively. The white block means a positive(+1) of occurrence state in the assemblage pattern.

From the Figures 5A, B and C, it can also be found that all the patterns were grouped into small clusters according to the LMP to which they were directed. These clusters can also be regarded as the ‘energy basins’ in the energy landscape, which indicate the pattern shifting tendency. Because of the corresponding relation between LMP and energy basin, there were four energy basins in the CD class and two energy basins in UC and non-IBD class. However, the component size was unbalanced in these energy basins; some basins were composed of a significant number of patterns while some were composed of only a few patterns: in the UC class, only 13 patterns were clustered to the basin with LMP 455, and the other 499 patterns belonged to the other basin with LMP 33. The additional figure 2 depicts the energy of each sample and the energy variation of each participant in ten time-series samples, which provides an overall illustration of the energy situation of the participants.

## 5 Discussion

### 5.1 The meaning of local minimal patterns

We constructed the energy landscape for an IBD data set by a pipeline incorporating the LDA model and the pairwise MaxEnt model. As defined by the pairwise MaxEnt model, the local minimal patterns(LMP) represents the system’s stage in high occurrence’ probability and high stability, in other words, the stable composition of the microbiota under a specific condition in our case.

### 5.2 Comparison of local minimal patterns between classes

Here we compared the LMPs between CD and healthy non-IBD classes to discover the alteration of microbiota when health conditions switch. Interestingly, three assemblages #1, #5 and #6 associated with the genus *Bacteroides* were observed as the only “activated” assemblage in three LMPs: P-#3, P-#17, and P-#33 of the CD class, respectively, while only the pattern P-#33 with activated assemblage A-#6 was in the non-IBD class. According to the probability of *Bacteroides*’ occurrence in assemblages ***φ***_*Bacter oides*_, *Bacteroides* genus was strongly dominant in the assemblage #6 with 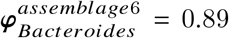, mildly dominant in the assemblage #1 with 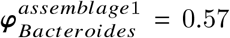 weakly dominant in the assemblage #5 with 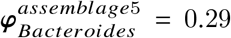 The decrease in the level of *Bacteroides* in CD and UC has been reported, and the level is even lower in an active phase than in remission[16]. The results we obtained may support the alternation of *Bacteroides* in the disease’ development of CD. Besides, the multiple LMPs characterized by different intensities of domination of *Bacteroides* may also suggest that the diversity of stable stages of microbiota in CD are affected by *Bacteroides*. If we consider the potential concurrence between the stage of disease’s development and microbiota, this result also implies the *Bacteroides*’s role as a potential marker of the disease.

Among these three patterns, genus *Alistipes* was the first dominant component in assemblage #5 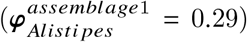 of LMP P-#17 apart from the *Bacteroides* in the other two patterns. *Alistipes* has been reported to relate to the gut inflammation but contrasting results about its contribution to the disease has also been reported[17]. Our result may support *Alistipes*’ harmful contribution to the CD’s development and this contribution might be be affected by the decrease of *Bacteroides*.

Interestingly, genus *Faecalibacterium* was the second dominant component in assemblage #1 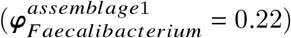 and assemblage #5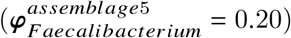 Note that the only species of this genus, *Faecalibacterium prausnitzii*, has been reported to decrease in the IBD pathogenesis[18] and have anti-inflammatory protein production[19]. Besides, Genus *Parabacteroides* was the third dominant member in assemblage #1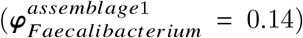 *Parabacteroides spp*. has been identified as probiotics and to be related to the alleviation of tumorigenesis and inflammations [20, 21]. Therefore, comparing between assemblages #6 and #1, the occurrence of activated assemblage #1 of CD can be interpreted as the joint effect of two positive factors and one negative factor: the increase of *Faecalibacterium*and *Parabacteroides*; the decrease of *Bacteroides*. We could speculate that the ‘trade-off’ between these factors from both directions and their contribution to CD’s development cause the LMP with activated assemblage #1 on the CD class.

Apart from three LMPs associated with *Bacteroides* of CD, in the last LMP P-#137 with activated assemblage #4 and assemblage #8, we can find the dominant genus of *Prevotella* and *Escherichia*, respectively. Both of them have been reported to be related to chronic inflammatory disease[22, 23]. Here we can suppose that the concurrence of *Prevotella* and *Escherichia* can be a potential feature of a particular stage in CD development.

We conclude that the genera mentioned above and the interaction of these genera might be the key to the alteration of microbiota in CD’s development, and how the potential ‘trade-off’ between them or other interaction modes contribute to the disease remain to be discovered. Besides, since the composition of an assemblage is complicated, further comprehensive understanding, especially the functional analysis from the metabolic aspect, is necessary.

### 5.3 How to consider the function of assemblage

All nine assemblages had their dominant components, and we may think these dominant genera can determine the assemblage’s contribution to the host’s microbiome to a great extent. Besides, we may also consider those genera which have a much higher probability to occur in one specific assemblage, or are ‘unique’ in a specific assemblage, will bring special features to the assemblage. However, as observed from the composition of the assemblages, most of the genera can satisfy the condition of ‘unique’, and this increases the complexity to studying the function of assemblages. Thus in this study we mainly discuss the function of assemblages according to their dominant components. The proper method in a more comprehensive and persuasive way is required to discover the assemblages in further studies.

### 5.4 The choice of assemblage number

According to the LDA model, every occurrence of a single taxon is assumed to belong to one specific latent character of ‘assemblage’ as what is defined in the model’s original application in natural language processing. Here the abundance data of microbiota were transformed into subcommunity, which we call microbial assemblage in this study. We chose the assemblage number of nine to achieve that every class has some relatively high abundance assemblages to control the loss of information during this transformation. The Figure 2A shows that, assemblages #3 and #8 had higher abundance in the CD class, assemblages #4, #7, and #9 had higher abundance in the UC class, and assemblage #2 was prominent in the non-IBD class. However, as mentioned in the result section, the assemblage abundance’s difference between classes was not significant and might have been caused by the fluctuation of a single assemblage in individuals; we can not simply conclude any single assemblage represents IBD or a healthy patient. And this may also apply to other diseases’ cases. Therefore, a comprehensive method considering the interaction between assemblages instead of considering them independently is necessary, which is also the purpose of applying the pairwise MaxEnt model in this study.

### 5.5 Effects of potential artifacts

In this study, two major artificial factors are introduced in our method. The first one is the choice of assemblage number as mentioned in the above section, which is nine in our case. Under the ideal condition, we can apply the pairwise MaxEnt modeling without assigning assemblage to each genus but by exploring all the pairwise interactions between the genera. However, the expensiveness with respect to computation is too high to finish the modeling for high dimensional input, and thus we introduced microbial assemblage as a dimensional reduction method. Because of the loss of information during the dimensional reduction, there is a trade-off between calculation performance and the ‘resolution’ of assemblage. Different choices of assemblage will change the modeling result and the interpretation of the result, so a number out of reasonable purpose and consideration is required.

Second is the threshold to define the occurrence state, which is 75 percentile of each class in our case. Because of the requirement of binarization of input data in the pairwise MaxEnt model, a threshold to define ‘relatively high’ is essential. The setting of the threshold will affect the modeling input as well as the modeling result. Here we do not have a precedent of guidelines or standards from previous studies or experiments. The 75 percentile is an intuitive definition of ‘relatively high’, but another threshold might also be considered and tested in our future’s studies.

### 5.6 The unbalance of energy basins’s size in energy basin

The patterns linked forward to the same LMP compose the clusters of patterns which we also call energy basins. We may consider the patterns within one energy basin to be similar and easy to transfer because each linked pair among them has an activity’s difference on only one assemblage and follows the steepest energy decline. Thus, speculation of the potential alteration direction of a specific pattern, or the stage of a microbial community, might also be implied from the basin. Here, a significant unbalance in basin’s size was observed in the UC class, where the basin characterized by LMP 455 only had 16 components patterns while the other basin characterized by LMP 33 had the other 496 component patterns. Here we suppose that the basin with a bigger size will have a more substantial effect since a random pattern might have a greater probability to be settled on the ‘track’ of this basin. However, a more thoughtful reason and meaning for the unbalance of basin’s size remain to be studied.

### 5.7 The value of the proposed method and its potential applications

Our proposed method integrates the Latent Dirichlet Allocation (LDA) topic model and the Pairwise Maximum Entropy (MaxEnt) model. The energy landscape quantifies all possible patterns of microbial community assembly and aids in elucidating the relationships between members under specific conditions. Although the energy landscape has previously been introduced for analyzing microbial data [24], it only considered the most abundant species. The LDA topic model makes it possible to consider the entire microbial community by clustering taxa into a few meaningful assemblages, serving as a dimension reduction. Our method removes the computational burden associated with the application of the Pairwise MaxEnt model to microbiome data and provides a valuable tool for time-series microbiome data analysis.

Furthermore, the energy landscape is constructed for a dynamic microbial community based on time-series microbiome data, which could provide insight into the alterations of the microbiota during disease progression and clarify the microorganism-host interaction. Its applications could be extended to explore the stability of microbial communities under specific health conditions and uncover the hallmarks of the microbiota that predispose hosts to develop diseases such as obesity, diabetes, colorectal cancer, and Alzheimer’s disease. This could contribute to the therapy and prognosis of these diseases.

In addition, the results of this method can be integrated with other biological tools to obtain a comprehensive understanding. For example, the metabolic characteristics of the microbial assemblages are crucial as the metabolic interactions within these assemblages likely drive the patterns of abundance. Combining these results with metabolism analysis tools, such as Flux Balance Analysis, could provide a more direct illustration of the relationship and even causality between the microbiota and disease.

## Supplementary information

**Supplementary Figure 1:**
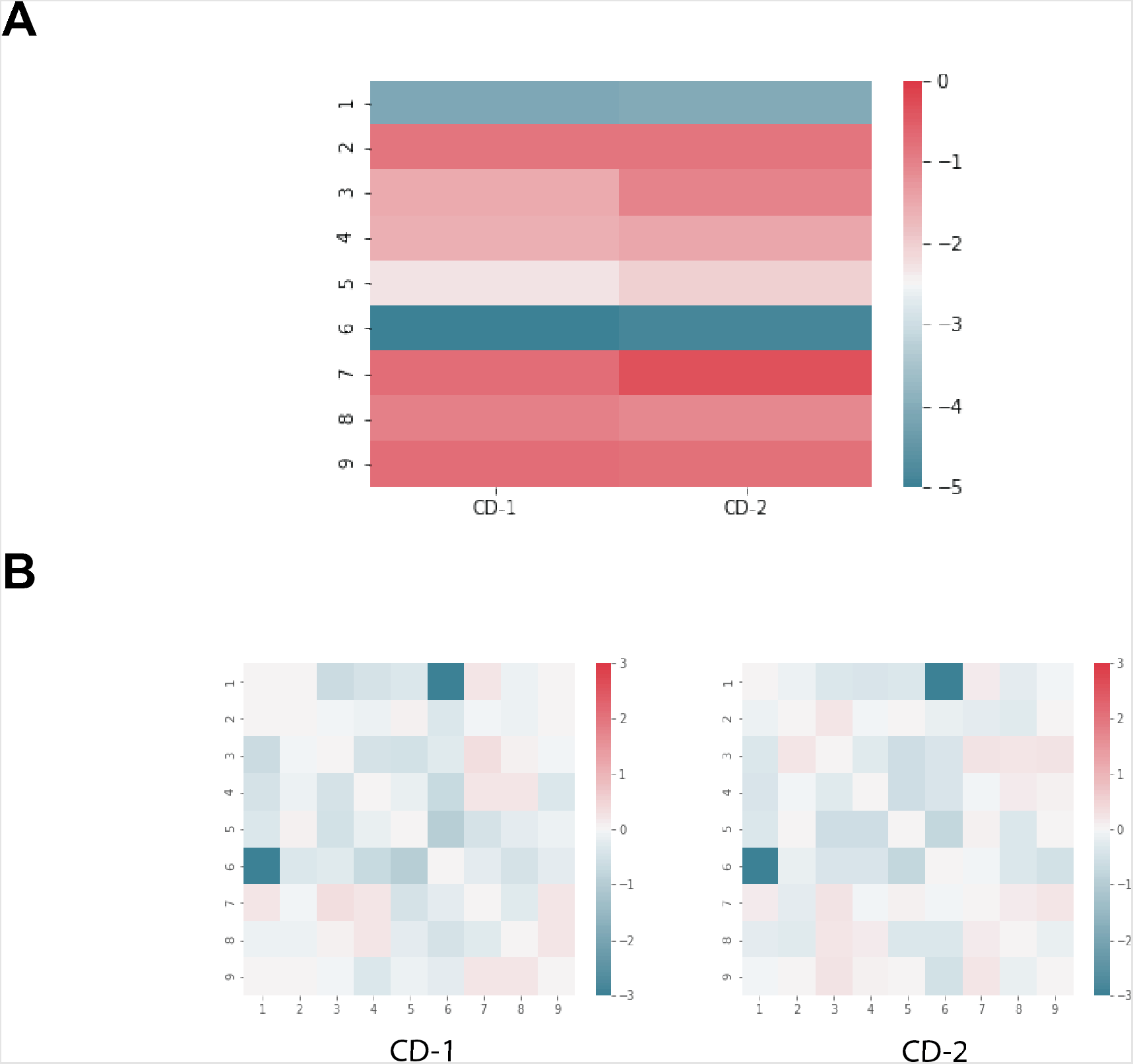
Application of Pairwise MaxEnt model for two different CD groups. **A**, the heatmap showing the tendency to occur of single assemblages ***h*** obtained from pairwise MaxEnt model. **B**, the heatmap showing pairwise interactions between assemblages ***g*** obtained from pairwise MaxEnt model.

**Supplementary Figure 2:**
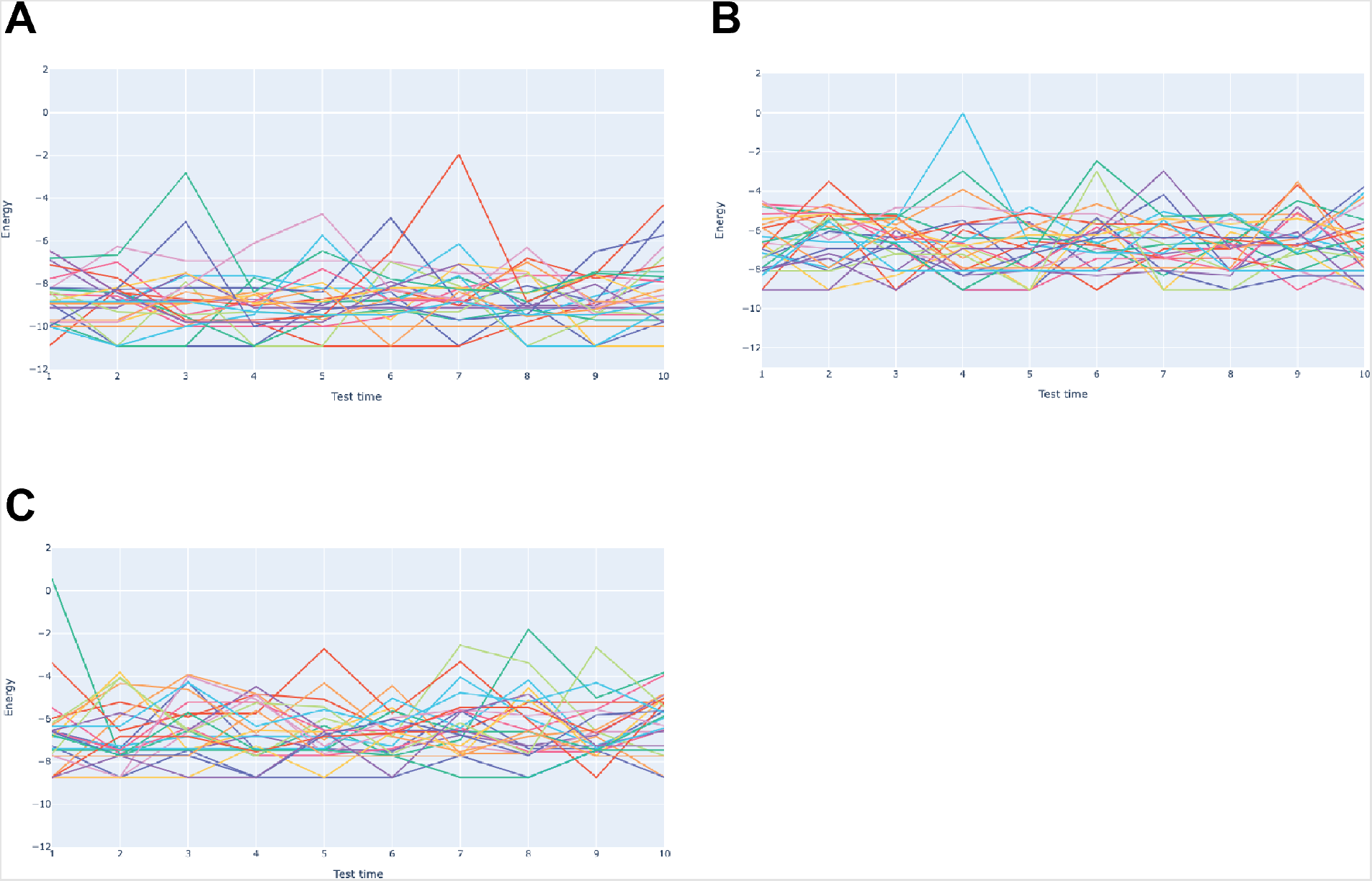
The energy variation of participants. **A, B, C** show the energy value assigned to all samples according to the modeling result in CD, UC, non-IBD classes, respectively. The lines represent single participants with ten time-series samples.

